# Inhibiting 5-lipoxygenase prevents skeletal muscle atrophy by targeting organogenesis signaling and insulin-like growth factor-1

**DOI:** 10.1101/2022.05.04.490621

**Authors:** Hyunjun Kim, Seon-Wook Kim, Sang-Hoon Lee, Da-Woon Jung, Darren R. Williams

## Abstract

**Background:** Skeletal muscle atrophy can occur in response to numerous factors, such as aging and certain medications, and produces a major socioeconomic burden. At present, there are no approved drugs for treating skeletal muscle atrophy. Arachidonate 5-lipoxygenase (Aox5) is a drug target for a number of diseases. However, pharmacological targeting of Alox5, and its role in skeletal muscle atrophy, is unclear.

**Methods:** The potential effects of gene knockdown and pharmacological targeting of Alox5 on skeletal muscle atrophy was investigated using cell-based models, animal models, and human skeletal muscle tissue cultures. Malotilate, a clinically safe drug developed for enhancing liver regeneration and Alox5 inhibitor, was investigated as a repurposing candidate. Mechanism(s) of action in skeletal muscle atrophy were assessed by measuring the expression level or activation status of key regulatory pathways, and validated using gene knockdown and RNA sequencing.

**Results:** Myotubes treated with the atrophy-inducing glucocorticoid, dexamethasone, were protected from catabolic responses by treatment with malotilate (+41.29%, P < 0.01). Similar anti-atrophy effects were achieved by gene knockdown of Alox5 (+30.4%, P < 0.05). Malotilate produced anti-atrophy effects without affecting the myogenic differentiation program. In an in vivo model of skeletal muscle atrophy, malotilate treatment enhanced muscle performance (Grip strength: +35.72%, Latency to fall: +553.1%, P < 0.05), increased mass and fiber cross sectional area (Quadriceps: +23.72%, Soleus: +33.3%, P < 0.01), and down-regulated atrogene expression (Atrogin-1: -61.58%, Murf-1: -66.06%, P < 0.01). Similar, beneficial effects of malotilate treatment were observed in an aging muscle, which also showed the preservation of fast twitch fibers (Type 2a: +56.48%, Type 2b: +37.32%, P < 0.01). Leukotrine B4, a product of Alox5 activity with inflammatory and catabolic functions, was found to be elevated in skeletal muscle undergoing atrophy (Quadriceps: +224.4%, P < 0.001). Cellular transcriptome analysis showed that targeting Alox5 upregulated biological processes regulating organogenesis and increased the expression of insulin-like growth factor-1, a key anti-atrophy hormone (+226.5%, P < 0.05). Interestingly, these effects were restricted to the atrophy condition and not observed in normal skeletal muscle cultures with Alox5 inhibition. Human skeletal muscle tissue was also protected from atrophy by pharmacological targeting of Alox5 (+23.68%, P < 0.05).

**Conclusion:** These results shed new light on novel drug targets and mechanisms underpinning skeletal muscle atrophy. Alox5 is a regulator and drug target for muscle atrophy, and malotilate is an attractive compound for repurposing studies to treat this disease.

## Introduction

Skeletal muscle atrophy causes muscle weakness that can progress to disability. It can be produced by a number of different factors, such as aging, certain medications, degenerative diseases or lifestyle factors [1]. Currently, the treatment options for skeletal muscle atrophy are limited to exercise and nutritional support, which suffer from variable outcomes and low compliance. Due to the significant economic impact produced by skeletal muscle atrophy and demographic aging in industrialized countries, there is a need to develop novel therapeutic options for treating this disorder [2].

Lipoxygenases are a family of non-heme, iron-containing enzymes that catalyze the dioxygenation of polyunsaturated fatty acids in lipids possessing a cis,cis-1,4-pentadiene hydrocarbon into molecules with diverse autocrine, paracrine, and endocrine signaling functions [3]. Arachidonate 5-lipoxygenase (Alox5; also known as 5-lipoxygenase, 5-LOX, or 5-LO), is one of the most widely studied lipoxygenases. Alox5 converts essential fatty acids into leukotrienes that possess a wide range of biological activities [4]. Consequently, Alox5 is under investigation as a drug target for numerous diseases [5]. However, the role of Alox5 in skeletal muscle atrophy is unclear. Endoplasmic reticulum (ER) stress was alleviated in myotubes after Alox5 inhibition and improved insulin resistance in a model of type 2 diabetes [6]. In contrast, genetic ablation of Alox5 failed to protect against muscle atrophy caused by denervation, whereas ablation of Alox12 or Alox15 was effective [7]. These results were supported by a report that baicalein, a phytochemical agent that also inhibits Alox12 and Alox15, maintained muscle fiber type and cross sectional area in ovariectomized rats [8]. Alox5 knockout in a model of post-menopausal osteoporosis produced increased skeletal muscle regenerative capacity after surgical injury, although this is distinct from the prevention of muscle fiber atrophy, which is characterized by dysregulated protein homeostasis and autophagy, and the preferential loss of particular myofiber types [9]. In addition, a recent study indicated that Alox5 gene expression was decreased, rather than upregulated, in a model of aging-induced sarcopenia [10].

Drug repurposing has significant advantages compared to traditional drug discovery approaches, because the repositioned compound is already characterized in other disease context(s) and clinical trials [11]. Malotilate (1,3-dithiol-2-ylidenepropanedioic acid bis(1-methylethyl) ester; CAS 59937-28-9) is a drug developed for treating liver disease [12]. It was shown to accelerate liver regeneration in rats, and inhibit inflammation and fibrosis [13]. Oral delivery in patients with advanced cirrhosis disease showed that malotilate was well tolerated with anti-fibrotic and hepatoprotective properties, and dry skin was the only possible adverse effect [14]. Clinical trials of malotilate treatment for primary biliary cirrhosis and alcoholic liver disease also reported small, beneficial effects [15]. Malotilate was shown to reduce the accumulation of type III and type IV collagen, laminin and fibronectin in a rat model of dimethylnitrosamine-induced liver fibrosis [16]. However, the first-pass elimination of malotilate was found to be dramatically reduced in cirrhotics, and a smaller amount of drug reached the liver in these patients [17]. Malotilate is included in the FDA-approved Drug Library (Selleck Chemicals, TX, USA) and is a specific inhibitor of Alox5, which regulates inflammatory responses [18]. The skeletal muscle of patients with pronounced aging-related atrophy, termed sarcopenia, is characterized by increased inflammation (known as ‘inflammaging’) and fibrosis [19]. In light of the therapeutic effects of malotilate treatment in studies of liver degeneration, inflammation and fibrosis, the potential for repurposing this drug for skeletal muscle atrophy was investigated.

In this study, the potential effects of pharmacological targeting of Alox5 on the progression of skeletal muscle atrophy was assessed by utilizing myotube-based models of skeletal muscle atrophy, animal models of muscle atrophy and aging, and human skeletal muscle tissue induced to undergo atrophy. The mechanism of action of Alox5 inhibition in skeletal muscle atrophy was assessed by measuring the expression level or activation status of major atrophy pathways, and validated using gene knockdown and cellular transcriptome analysis.

## Materials and methods

### Reagents

Dexamethasone (Dex) was purchased from Santa Cruz Biotechnology, TX, USA. Malotilate was purchased from Selleckchem, TX, USA. Leukotriene B4 (LTB4) was purchased from Tocris, Bristol, UK.

### Cell culture

C2C12 murine skeletal muscle myoblasts were purchased from Koram Biotech. Corp, Republic of Korea. The cells were cultured with growth media (GM), consisting of Dulbecco’s Modified Eagle’s Medium (DMEM), 10% fetal bovine serum (FBS), and 1% penicillin and streptomycin (PenStrep). Myoblasts were differentiated into myotubes at approximately 90% confluence, by culturing in differentiation media (DM: DMEM, 2% horse serum (HS) and PenStrep) for 96 h. Myotubes were treated with 10 μM Dex for 24 h to induce atrophy. To assess myogenesis, differentiating myoblasts were treated with compound of interest at the same time as culture in DM for 24 h or 48 h.

### Immunocytochemistry

Myotubes were visualized by myosin heavy chain immunocytochemistry. In brief, myotubes were washed three times with PBS and fixed with 3.7% formaldehyde solution, washed with PBS, and permeabilized with 0.2% Triton X-100 solution. After further washing with PBS, the myotubes were blocked with 1% bovine serum albumin in PBST (0.1% Tween 20 in PBS) and incubated with the primary antibody overnight at 4 °C, followed by washing with PBS and treatment with the secondary antibody for 1 h. The myotubes were then counterstained with 1 µM of DAPI in PBS. Fluorescence images of stained myotubes were obtained in 5 different areas using a DMI 3000 B microscope (Leica, Germany) and analyzed using ImageJ 1.52 software (National institutes of Health, Bethesda, MD, USA). Myotubes were classified as myosin heavy chain positive cells containing more than 4 nuclei.

### Western blotting

Cell protein lysates were quantified using the Bradford reagent (Bio-Rad, USA, CA) and separated by electrophoresis with 10% or 12% SDS-PAGEs. The separated proteins were transferred onto PVDF membranes (Merck, Germany), blocked with 5% non-fat powdered milk in 0.02% Tween 20 in TBS (TBST) and 5% bovine serum albumin in TBST, and incubated overnight at 4°C with the primary antibody of interest. The secondary antibody was used at a 1: 2000 dilution, with incubation for 35 min at RT. Densitometry analysis was carried out using ImageJ 1.48 software (National Institutes of Health, Bethesda). Details of the primary and secondary antibodies are provided in Supplementary Table 1 and Table 2, respectively.

### Real-time qPCR

mRNA levels of the genes of interest was measured using the StepOnePlus Real Time PCR System (Applied Biosystems, UK). cDNA was synthesized from total RNA using the AccuPower RT PreMix (Bioneer, Republic of Korea). Real-time PCR (qPCR) was conducted according to the manufacturer’s instructions with the following modifications: PCR was performed in triplicate in 20 μL 2X Power SYBR Green PCR Master Mix (Enzynomics, Republic of Korea) composed of 200 nM (final concentration) of the targeted primers and 1 μL of cDNA. Details of the primers used in this study are provided in Supplementary Table 3. The denaturation step was conducted by incubation for 10 min at 95 °C and the amplification step consisted of 40 cycles of denaturation (15 s at 95 °C), annealing (1 m at 60 °C) and extension (72 °C for 20 s). The fluorescence at the extension step was detected at 72 °C after each cycle. After the final cycle, the melting-point of the samples was analyzed within the range of 60–95 °C using fluorescence detection. A specific cDNA sample was included in each run and compared as a reference between runs.

The expression level of β-actin was used for normalization while measuring the expression levels of all of the other genes.

### LTB4 ELISA

An LTB4 ELISA kit was purchased from My Biosource, CA, USA. The concentrations of LTB4 in skeletal muscles and cell lysates were measured by ELISA according to the manufacturer’s instructions. Samples (100 μL) were incubated with polyclonal LTB4 binding in 96 wells for 1 h. After stopping the enzymatic reaction, optical density was measured at 405 nm.

### Animal studies

Animal studies were carried out under the auspices of the Institute for Laboratory Animal Research Guide for the Care and Use of Laboratory Animals, and approved by the Gwangju Institute of Science and Technology Animal Care and Use Committee (study approval number GIST-2021-059). Animal studies have been approved by the appropriate ethics committee and have therefore been performed in accordance with the ethical standards laid down in the 1964 Declaration of Helsinki and its later amendments. Animals were purchased from Damool Science, Republic of Korea.

### Dexamethasone model of skeletal muscle atrophy

12 weeks old male C57BL/6J mice were treated with drugs as follows: 1) Injection of vehicle (5% DMSO, 5% Tween80, PBS) alone, 2) 15 mg/kg dexamethasone dissolved in vehicle, 3) Injection of 15 mg/kg dexamethasone and 10 mg/kg malotilate, (n=5 per group). Mice were treated by intraperitoneal injection every 24 h for 14 d and then assessed for skeletal muscle function and condition.

### Aging model of skeletal muscle atrophy

20 month old male C57BL/6J mice were treated with drugs as follows: 1) Vehicle alone (6.73% DMSO in 0.5% carboxyl methyl mellulose), 2) 50 mg/kg malotilate, (n=5 per group). Mice were treated by oral gavage every 24 h for 4 weeks and then assessed for skeletal muscle function and condition.

### Measurement of grip strength

Grip strength was measured with the BIO-GS3 grip strength test meter (Bioseb, FL, USA). Mice were placed onto the grid with all four paws attached and gently pulled backwards to measure grip strength until the grid was released. The maximum grip value used to represent muscle force was obtained using 3 trials with a 30 second interval.

### Muscle fatigue test

Muscle fatigue was measured with two different models by using the Rotarod. One is the constant model and the other is the accelerating model that is inherent in the rotarod machine. In brief, the mice were accommodated to training before starting the fatigue task using an accelerating rotarod (Ugo Basile, Italy). Mice were trained with speeds ramping from 10 rpm to 15 rpm. 24 h later, the muscle fatigue test was carried out rotarod running with 5 rpm increments every 5 min up to 15rpm. Latency to fall off the rotarod for each mouse was measured. A fatigued mouse was classified as falling off 4 times within 1 min, which then terminated the test. Maximum rpm was measured by acceleration model inherent in the machine and recorded the rpm which the mice fell off from the machine.

### Muscle dissection and histological analysis

Mice were anesthetized with 22 mg/kg ketamine (Yuhan, Republic of Korea) and 10 mg/kg xylazine (Bayer, Republic of Korea) by IP injection before sacrifice. Quadriceps, gastrocnemius, tibialis anterior and soleus muscles were dissected and weighed. For immunohistochemistry, the muscles were then fixed by overnight incubation with 4% paraformaldehyde at 4 °C, embedded into paraffin solution and stored -80 °C. Sections were obtained using a cryostat. Sectioning and hematoxylin and eosin (H&E) staining was carried out by the Animal Research Facility of the Gwangju Institute of Science and Technology, Republic of Korea. H&E staining was carried out using a kit (Merck, Germany). Muscle fiber cross sectional area was measured with the ImageJ 1.48 software (National Institutes of Health, USA).

### Immunohistochemistry

Immunohistochemistry was carried out using anti-myosin heavy chain type 2A, and 2B antibodies (DSHB, IA, USA) and an anti-laminin antibody (Abcam, Cambirdge, UK). Counterstaining was conducted by using 1 μM of DAPI solution. Muscle fiber cross sectional area was measured with the ImageJ 1.48 software (National Institutes of Health, USA) after the images were visualized with fluorescence microscopy (LEICA DM 2500).

### RNA Seq

RNA samples were obtained from C2C12 murine myoblasts cultured as follows (1) differentiation media (DM) for 120 h (Un); (2) DM for 96 h and DM plus 10 μM Malotilate for 24 hours (Mal); (3) DM for 96 h and DM plus 10 μM Dex for 24 h (Dex); (4) DM for 96 h and DM plus 10 μM Dex and 10 μM malotilate for 24 h (Dex_Mal); RNA Seq was carried out by Macrogen, Republic of Korea. Before proceeding sequencing, QC was performed by FastQC v0.11.7(http://www.bioinformatics.babraham.ac.uk/projects/fastqc/) . Illumina paired-ends or single ends in sequenced samples were trimmed by Trimmomatic 0.38(http://www.usadellab.org/cms/?page=trimmomatic) with various parameters. Sequences of samples were analyzed and mapped by HISAT2 version 2.1,0, Bowtie2 2.3.4.1(https://ccb.jhu.edu/software/hisat2/index.shtml). Potential transcripts and multiple splice variants were assembled by StringTie version 2.1.3b(https://ccb.jhu.edu/software/stringtie/). 15619 genes were quantified in total. Heat map in Figure 7F) was made by Graphpad Prism 7 (Graphpad, CA, USA).

### Human skeletal myoblasts culture and experimentation

Human skeletal myoblasts were obtained from Thermo-Fisher Scientific, USA). The myoblasts were thawed using a water bath, centrifuged at 180 g for 5 min at room temperature washed with 20 mL DM. Myoblasts were then re-suspended in DM and seeded onto 12 well plates at a density of 4.8 x 10^4^ cells/well. After 48 h later, myoblasts were treated with compounds of interest for 24 h. Myotubes were stained with H&E. Myotube diameter was measured using the ImageJ 1.48 software (National Institutes of Health, USA) after light microscopy analysis of DIC captured images (Olympus CKX41)

### Statistical analysis

Statistical significance was determined using The Student’s *t*-test. A *p* value of less than 0.05 was considered as significant. Unless otherwise mentioned, all data shown are representative of more than 3 experimental repeats and the graph error bars are standard deviation.

## Results

### Malotilate inhibited atrophy in glucocorticoid-treated murine myotubes

The glucocorticoid treatment model was used to investigate the effect of malotilate on myotube atrophy. This model was selected because glucocorticoid levels are increased during aging and multiple types of skeletal muscle atrophy, such as sepsis, starvation, chronic obstructive pulmonary disease, diabetes and cancer. Murine myotubes treated with the glucocorticoid, Dex, showed reduced diameter and a larger proportion of thinner myotubes, which indicate myotube atrophy. Malotilate treatment inhibited the effects of Dex on myotube atrophy (Figure 1A-B). Atrogin-1 and MuRF-1 are important atrogenes that become upregulated in skeletal muscle wasting. Malotilate treatment inhibited the upregulation of Atrogin-1 and MuRF-1 in Dex treated myotubes (Figure 1C-D). Moreover, total protein synthesis was decreased by treatment with Dex and recovered by treatment with malotilate (Figure 1E-F).

**Figure 1:**
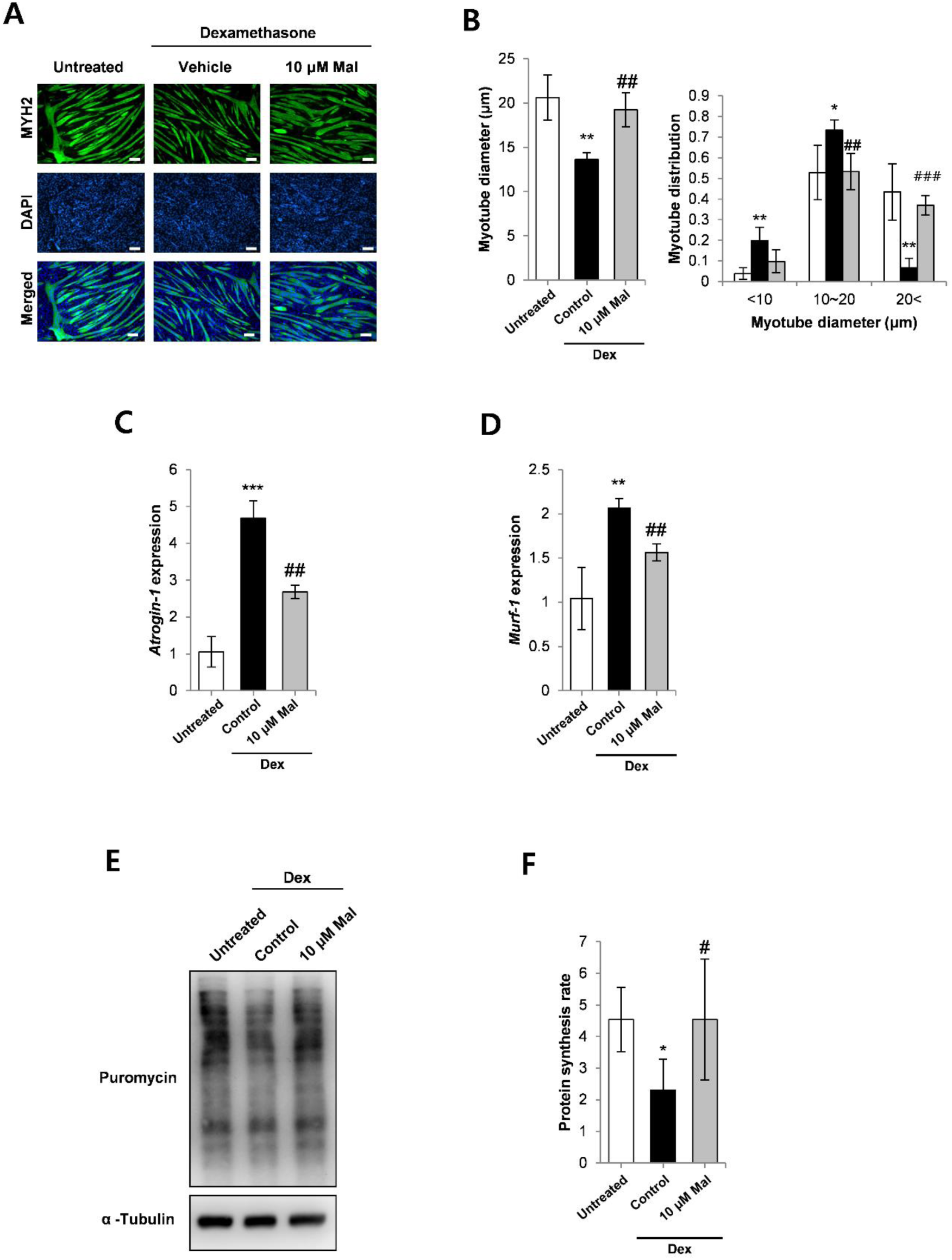
A) Myosin heavy chain (MYH2) immunocytochemistry of C2C12 myoblasts cultured as follows: (1) differentiation media (DM) for 120 h (untreated); (2) DM for 96 h and DM plus 10 μM Dex for 24 hours; (3) DM for 96 h and DM plus 10 μM Dex and 10 μM malotilate (Mal) for 24 h (scale bar=100 μm). B) Myotube diameter and myotube diameter distribution. C-D) qPCR analysis of atrogin-1 and MuRF-1 expression. E-F) SUnSET assay of protein synthesis. α-Tubulin expression was used as the loading control. *=*p*<0.05, **=*p*<0.01, ***=*p*<0.001 compared to untreated. ^#^=*p*<0.05, ^##^=*p*<0.01, ^###^=*p*<0.001 compared to Dex treated.

The transcription factor forkhead box O3 (FoxO3a) is a major regulator of the ubiquitin-proteasome pathway of skeletal muscle atrophy. De-phosphorylation of FoxO3a by catabolic signaling pathways induces its translocation from the cytoplasm to the nucleus and upregulation of target atrogenes. Foxo3a phosphorylation levels were decreased in myotubes treated with Dex and increased by treatment with malotilate (Figure 2A-C), and accompanied by a decrease in atrogin-1 expression (Figure 2D). Dysregulated autophagy is a key feature of skeletal muscle atrophy. Dex treatment increased autophagy in the myotubes (increased ratio of LC3bII to LC3bI, and reduced expression of p62). Autophagy in the Dex treated myotubes was decreased by treatment with malotilate (Figure 2E-F).

**Figure 2:**
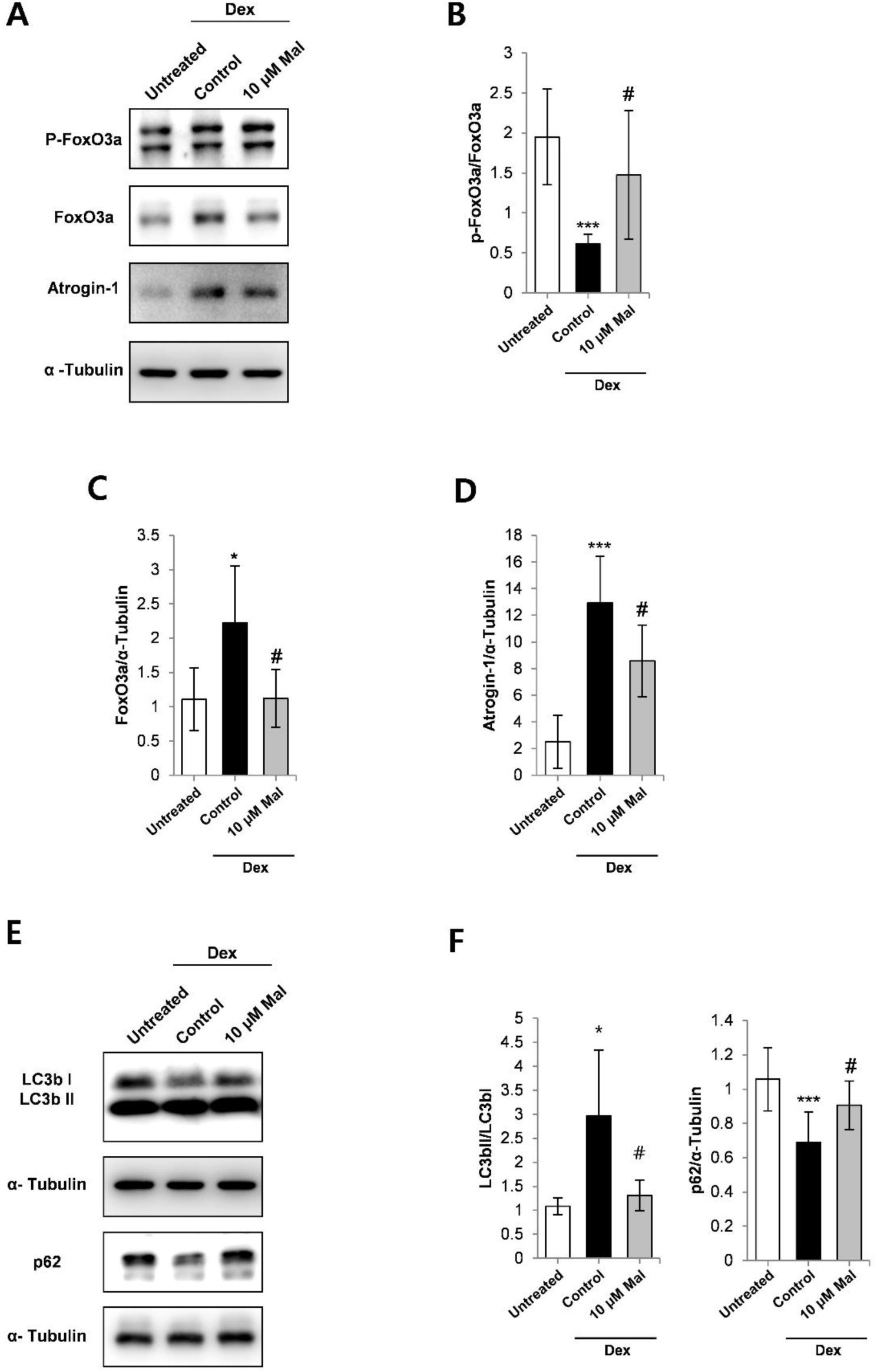
A-D) Western blot analysis of FoxO3a phosphorylation, FoxO3a expression, and atrogin-1 expression. α-Tubulin expression was used as the loading control. E-F) Western blot analysis of LC3b I, LC3b II, and p62 expression. α-Tubulin expression was used as the loading control. *=*p*<0.05, ***=*p*<0.001 compared to untreated. ^#^=*p*<0.05 compared to Dex treated.

### Alox5 gene knockdown prevents myotube atrophy associated with increased levels of the Alox5 product, leukotriene B4

Malotilate has been characterized as a selective inhibitor of Alox5. The effect of Alox5 gene knockdown on myotube atrophy was assessed in Dex treated myotubes. The ability of siRNA to reduce Alox5 expression was confirmed by qPCR and western blotting (Figure 3A-B). Alox5 gene knockdown inhibited myotube atrophy (Figure 3C-D). Gene knockdown was accompanied by a reduction in atrogin-1 expression (Figure 3E). Treatment with the Alox5 enzyme inhibitor, malotilate, did not affect Alox5 expression in Dex-treated myotubes (Figure 3F).

**Figure 3:**
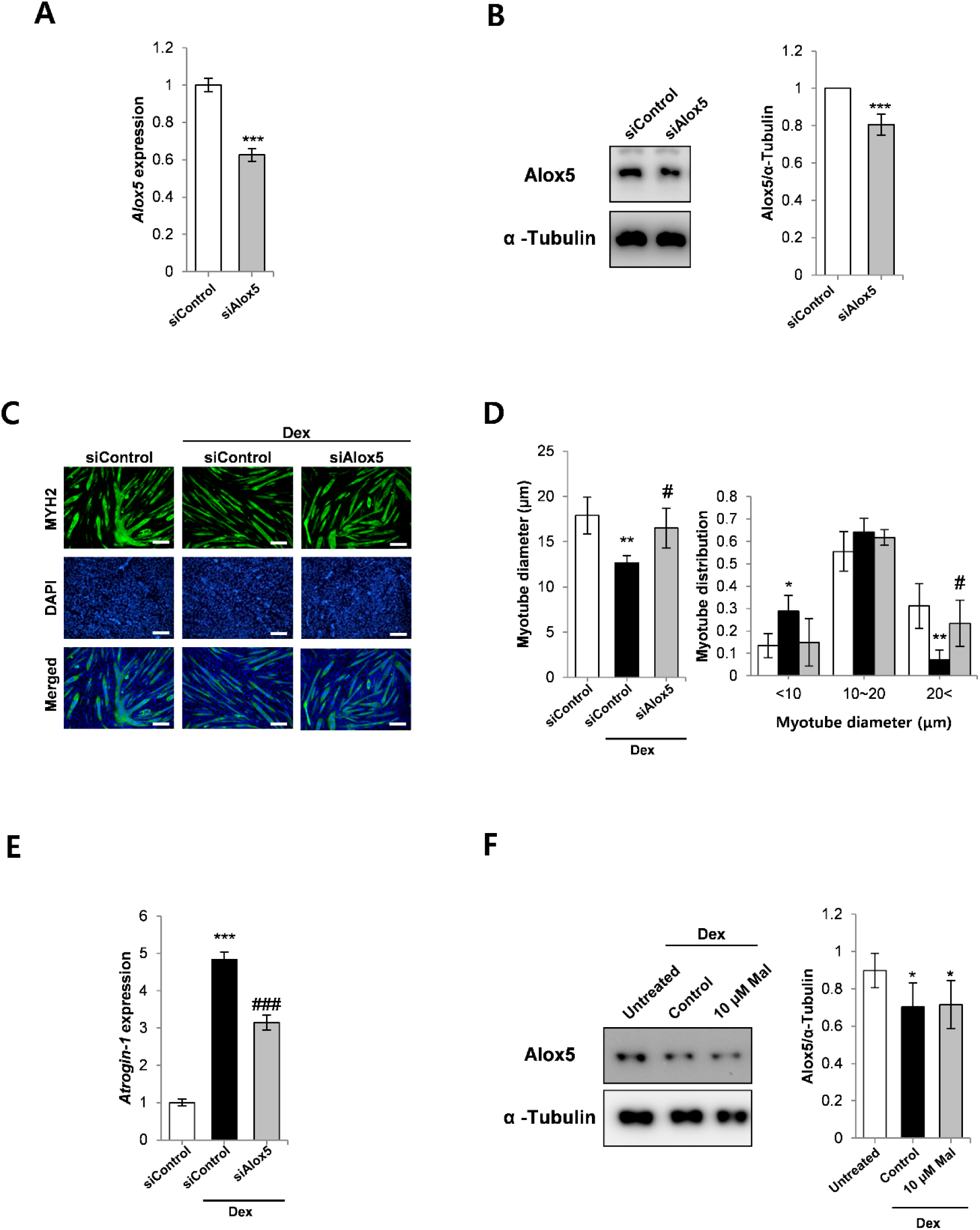
A) qPCR analysis of Alox5 expression in C2C12 myoblasts cultured as follows: (1) 72 h incubation with DM and 48 h incubation with scrambled siRNA; (2) 72 h incubation with DM and 48 h incubation with Alox5 siRNA. B) Western blot analysis of Alox5 expression in C2C12 myoblasts cultured as described in part (A). C) MYH2 immunocytochemistry of C2C12 myoblasts cultured as follows: (1) 72 h incubation with DM and 48 h incubation with DM plus control, scrambled siRNA; (2) Following 72 h incubation with DM, 24 h incubation in with DM plus control, scrambled siRNA and additional 24 h treatment with 10 μM Dex plus scrambled siRNA; (3) Following 72 h incubation with DM, 24 h incubation in with DM plus control, scrambled siRNA and additional 24 hour treatment with 10 μM Dex plus Alox5 siRNA; (scale bar=100 μm). D) Myotube diameter and distribution. E) qPCR analysis of atrogin-1 expression in C2C12 myoblasts cultured as described in part (C). F) Western blot analysis of Alox5 expression in C2C12 myoblasts cultured as follows: 1) 120 h incubation with DM; (2) 96 h incubation with DM, and 24 hour treatment with 10 μM Dex; (3) 96 h incubation with DM, and 24 hour treatment with 10 μM Dex and 10 μM malotilate (Mal). For (A-B, F): *=*p*<0.05, ***=*p*<0.001 compared to untreated. ^#^=*p*<0.05, ^###^=*p*<0.001 compared to Dex treatment alone. For (D-E): *=*p*<0.05, **=*p*<0.01, ***=*p*<0.001 compared to control siRNA. ^#^=*p*<0.05, ^###^=*p*<0.001 compared to control siRNA plus Dex.

Leukotriene B4 (LTB4) is an Alox5 product that is involved in inflammatory responses and catabolic pathways, such as NF-κb and FoxO signaling. LTB4 levels were shown to be increased in Dex-treated myotubes and lowered by malotilate (Supplementary Figure 1A). Accordingly, application of exogenous LTB4 did not improve atrophy in Dex-treated myotubes or reduce the expression of atrogin-1 (Supplementary Figure 1B-D). To confirm that drug-mediated inhibition of Alox5 prevented atrophy, myotubes were treated with SPI-1005, which is a small molecule inhibitor of Alox5. SPI-1005 treatment reduced LTB4 levels in myotubes (Supplementary Figure 1E), prevented Dex-induced atrophy (Supplementary Figure 1F-G), and inhibited the induction of atrogin-1 expression (Supplementary Figure 1H).

### Malotilate does not significantly affect the myogenic program

To assess the effect of malotilate treatment on myogenesis, murine myotubes were induced to differentiate into myotubes in the presence of malotilate. The expression of myosin heavy chain type 2 (Myh2), a marker of differentiation, was not affected by malotilate treatment during myogenesis (Supplementary Figure 2A-B). Increased phosphorylation of Akt is a key signaling event during myogenesis. In differentiated myotubes, the phosphorylation level of Akt was not significantly changed by malotilate treatment (Supplementary Figure 2C). Expression analysis of master genetic regulators the myogenic program (Pax7, Myf5, MyoD and MyoG[20]) did not show any significant change after 24 h malotilate treatment (Supplementary Figure 2D).

### Pharmacological targeting of Alox5 inhibits skeletal muscle atrophy in a glucocorticoid treatment model

As first test to investigate the effectiveness of malotilate to prevent skeletal muscle atrophy *in vivo*, the Dex glucocorticoid treatment model was selected. Malotilate treatment for 14 d did not significantly affect body weight in the Dex treated mice (Supplementary Figure 3A-B). Malotilate treatment significantly increased grip strength in the Dex treated mice and reduced latency to fall in the rotarod performance test (Figure 4A). Maximum running speed was also significantly increased by malotilate treatment (Supplementary Figure 3C). ELISA analysis indicated that levels of the Alox5 product, LTB4, were increased in Dex-treated muscle and lowered by malotilate treatment (Figure 4B). Mean muscle fiber cross sectional area (CSA) and the proportion of fibers with larger CSA was increased by malotilate treatment (Figure 4C-D and Supplementary Figure 3D). Assessment of skeletal muscle mass indicated that malotilate treatment increased mass in the quadriceps and soleus muscles of dexamethasone treated mice (Supplementary Figure 3E). Malotilate treatment also decreased expression of the atrogenes, atrogin-1 and MuRF-1, in the Dex treated mice (Figure 4E-F).

**Figure 4:**
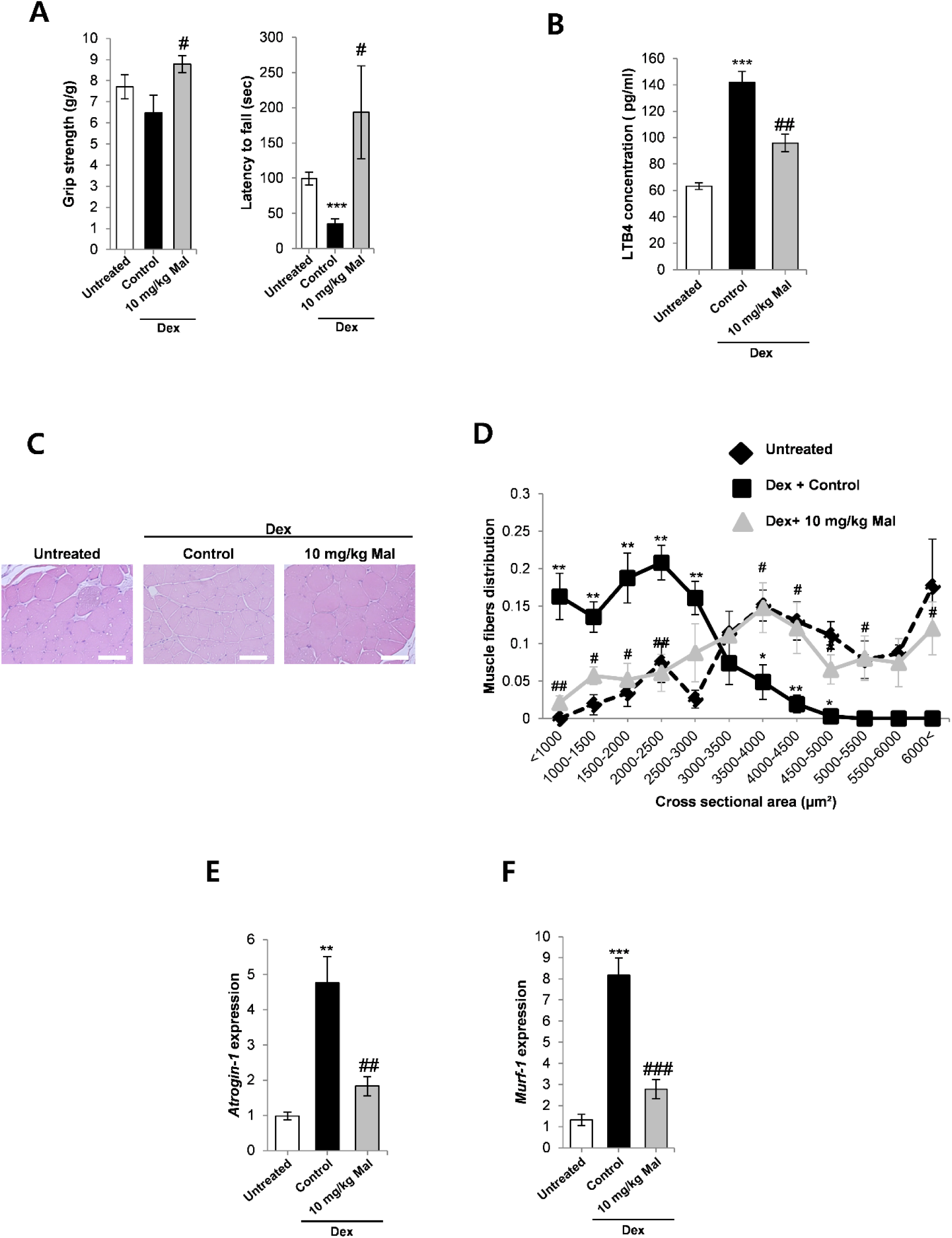
A) Grip strength and latency to fall off the rotarod in mice treated with vehicle alone, Dex, or Dex plus malotilate (Mal) for 14 d. B) ELISA-based quantification of the Alox5 product, LTB4, in the quadriceps. C) Representative H&E staining of the quadriceps (scale bar=100 μm). D) Fiber cross sectional area in the quadriceps. E-F) qPCR analysis of atrogin-1 and MuRF-1 expression. *=*p*<0.05, **=*p*<0.01, ***=*p*<0.001 compared to vehicle treated mice. ^#^=*p*<0.05, ^##^=*p*<0.01, ^###^=*p*<0.001 compared to Dex treated mice (n=5 per experimental group).

### Pharmacological targeting of Alox5 inhibits skeletal muscle atrophy in aged mice

To assess the potential of malotilate to treat aging-related skeletal muscle atrophy (termed sarcopenia), aged mice were treated by oral delivery for 4 weeks. Malotilate treatment did not affect body mass in the aged mice (Figure 5A). Grip strength was significantly increased by malotilate treatment (Figure 5B). The mean latency to fall was increased in the malotilate treated mice, but did not reach statistical significance (Figure 5C). Histological analysis of the quadriceps muscle showed that average myofiber cross sectional area was increased by malotilate treatment (Figure 5D-E). Myofiber diameter distribution indicated a significantly greater proportion of large myofibers (>5000 μm^2^) in malotilate treated aged mice (Figure 5F). Quadriceps and soleus muscle mass were significantly increased in malotilate treated aged mice (Figure 6A-B). Immunohistological analysis of fast twitch fiber types IIA and IIB, which preferentially undergo atrophy during aging compared to slow class I twitch fibers [21], indicated that malotilate treatment preserved the CSA of fast fibers (Figure 6C-F).

**Figure 5:**
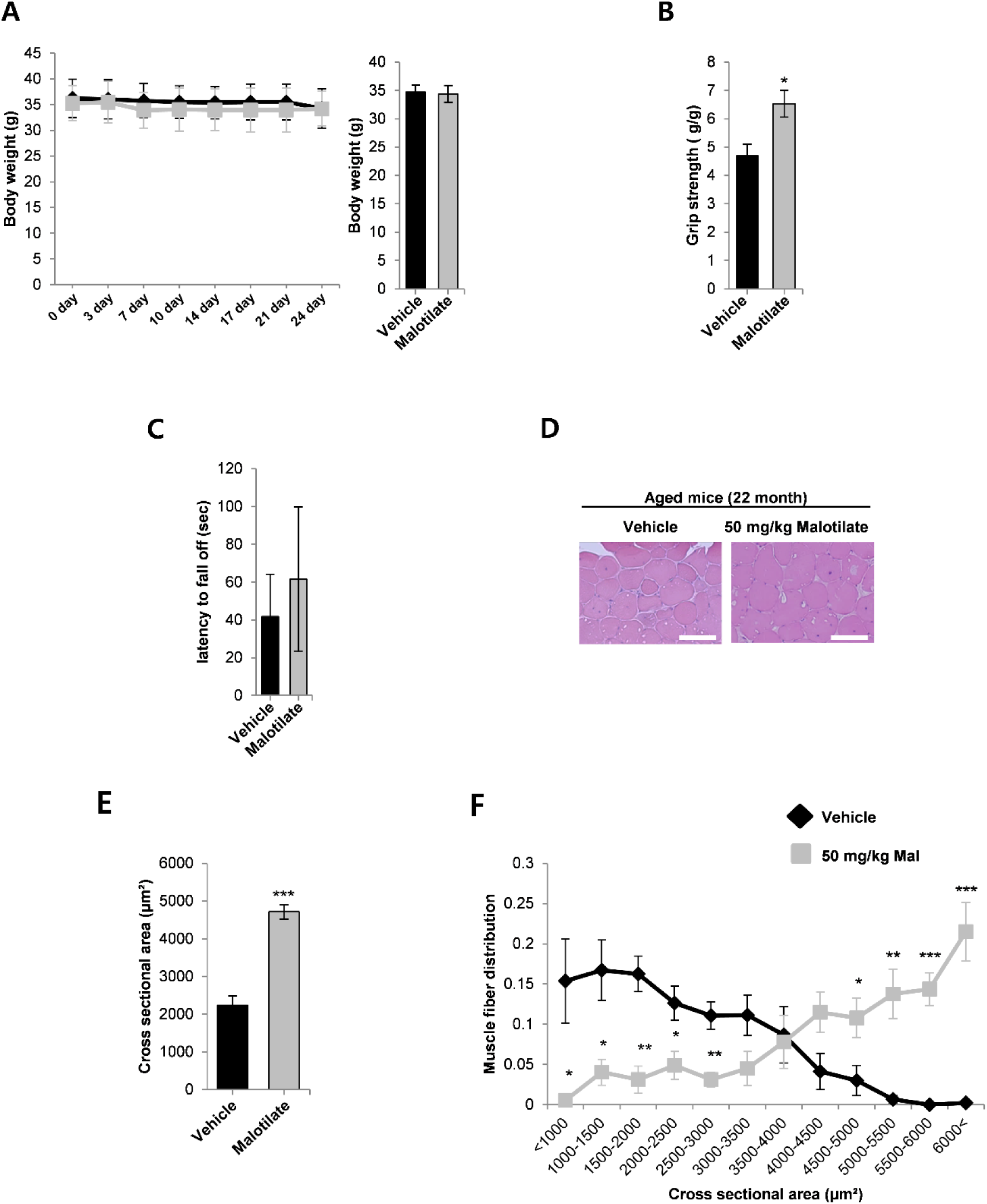
A) Body weight and final body weight in aged (22 months) after 4 weeks treatment with vehicle or malotilate (Mal). B) Grip strength. C) Latency to fall off the rotarod. D) Representative H&E staining of the quadriceps (scale bar=100 μm). E-F) Cross sectional fiber area and muscle fiber distribution of the quadriceps muscle. *=*p*<0.05, ***=*p*<0.01, ***=*p*<0.001 compared to vehicle treated (n=5 per experimental group).

**Figure 6:**
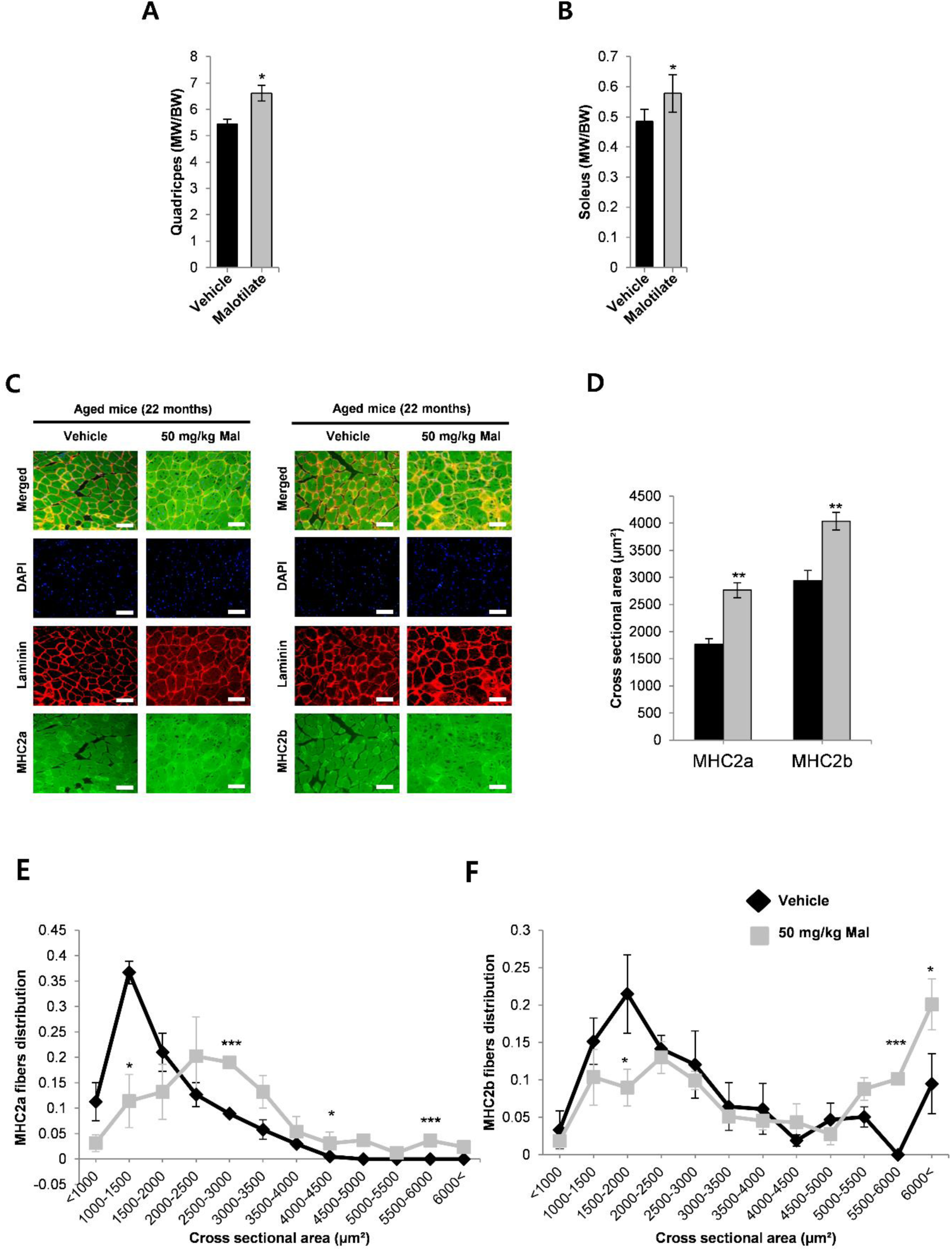
A) Quadriceps muscle mass in aged (22 months) after 4 weeks treatment with vehicle or malotilate (Mal). B) Soleus muscle mass. C) Representative images of myosin type 2a and 2b, and laminin immunostaning in the quadriceps muscle. DAPI was used to visualize cell nuclei (scale bar=100 μm). D) Mean cross sectional area of type 2a and type 2b fibers in the quadriceps. E-F) Cross sectional area distribution of type 2a and type 2b fibers in the quadriceps. *=*p*<0.05, ***=*p*<0.01, ***=*p*<0.001 compared to vehicle treated (n=5 per experimental group).

**Figure 7:**
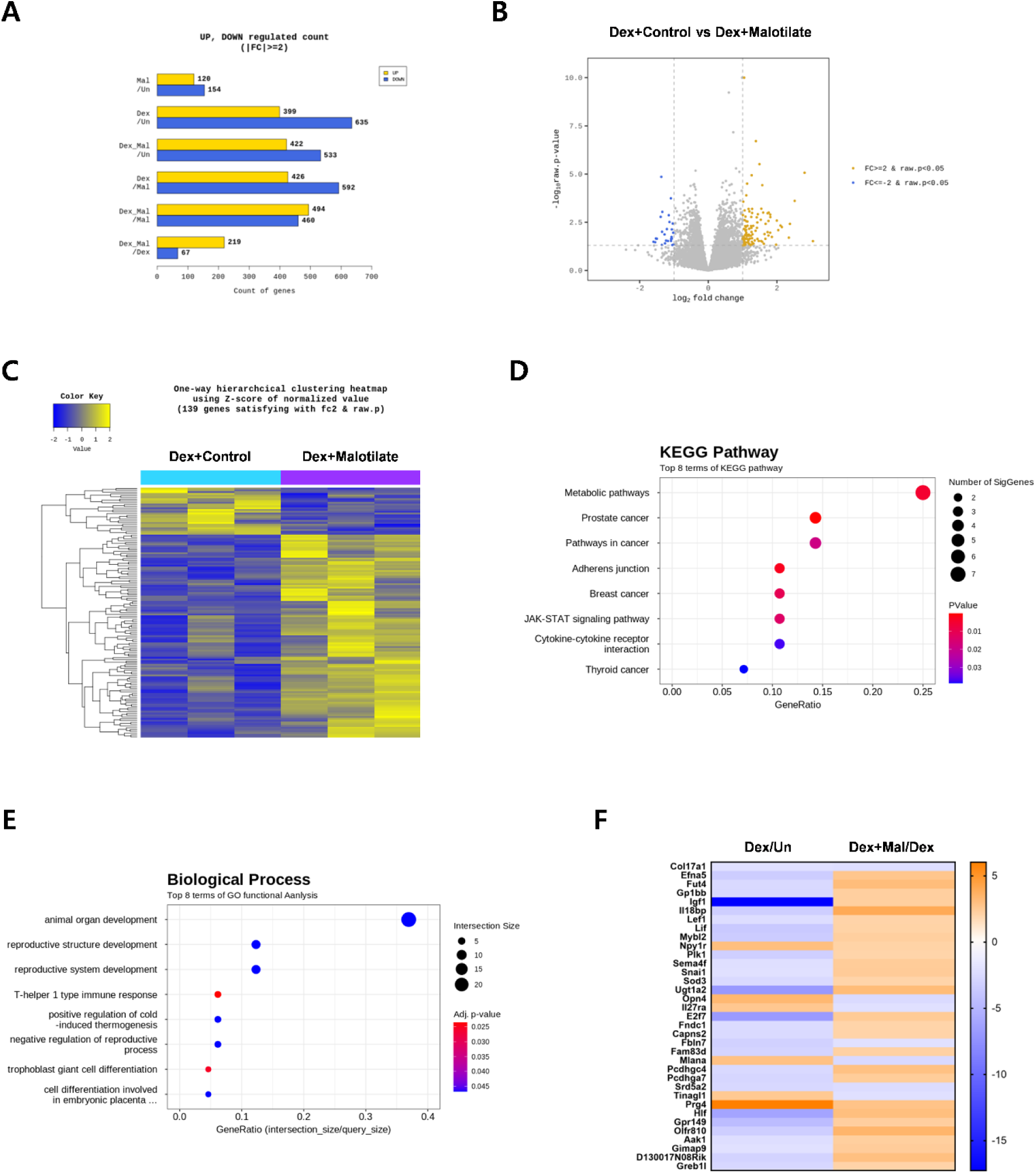
A) Cellular transcriptome analysis showing total number of upregulated and downregulated genes between C2C12 murine myoblasts cultured as follows (1) differentiation media (DM) for 120 h (Un); (2) DM for 96 h and DM plus 10 μM Malotilate for 24 hours (Mal); (3) DM for 96 h and DM plus 10 μM Dex for 24 h (Dex); (4) DM for 96 h and DM plus 10 μM Dex and 10 μM malotilate for 24 h (Dex_Mal). B) Volcano plot showing gene expression changes in the Dex compared to Dex_Mal treatment groups. C) One way hierarchical heat map for the Dex compared to Dex_Mal treatment groups. D) Gene ontology (GO) functional analysis for the Dex compared to Dex_Mal treatment groups. E) KEGG (Kyoto Encyclopedia of Genes and Genomes) pathway analysis for the Dex compared to Dex_Mal treatment groups. F) Heat map for genes showing differential expression in Dex and Dex_Mal, compared to Mal and Un.

### Cellular transcriptomic analysis of myotubes with pharmacological targeting of Alox5 in the presence or absence of atrophy

To gain an overview of the mechanism by which Alox5 inhibition by malotilate inhibits muscle atrophy, myotubes undergoing Dex-induced atrophy were treated with malotilate and the cellular transcriptome was investigated using RNA Seq. Expression patterns were compared with untreated Dex treated myotubes, normal myotubes or myotubes treated with malotilate alone (Figure 7A). The heat map and volcano plots showed a total of 139 differentially expressed genes (DEGs) between myotubes treated with Dex or Dex plus malotilate (Figure 7B-C). KEGG (Kyoto Encyclopedia of Genes and Genomes) pathway analysis showed that metabolic pathways were primarily targeted by malotilate treatment (Figure 7D). Gene ontology (GO) functional analysis indicated that organ development was the top biological process targeted in myotubes treated with Dex plus malotilate compared to Dex alone (Figure 7E). A heat map for genes showing differential expression in Dex plus malotilate and Dex alone, compared to malotilate and untreated, is shown in Figure 7F. The expression of a number of genes known to be linked to skeletal muscle atrophy were regulated by malotilate treatment. Genes which are known to be linked to skeletal muscle atrophy are shown in Supplementary Figure 4.

### Pharmacological targeting of Alox5 prevents atrophy in human skeletal muscle cultures

Skeletal muscle precursor cells from human donors were differentiated into myotubes and induced to undergo atrophy by treatment with dexamethasone. Histological analysis indicated that co-treatment with malotilate could prevent atrophy in the human myotubes, as shown by increased average myotube diameter and greater proportion of larger sized myotubes (Figure 8A-C). Malotilate treatment also significantly reduced expression of the atrogenes, atrogin-1 and MuRF-1, in the human myotubes (Figure 8D).

**Figure 8:**
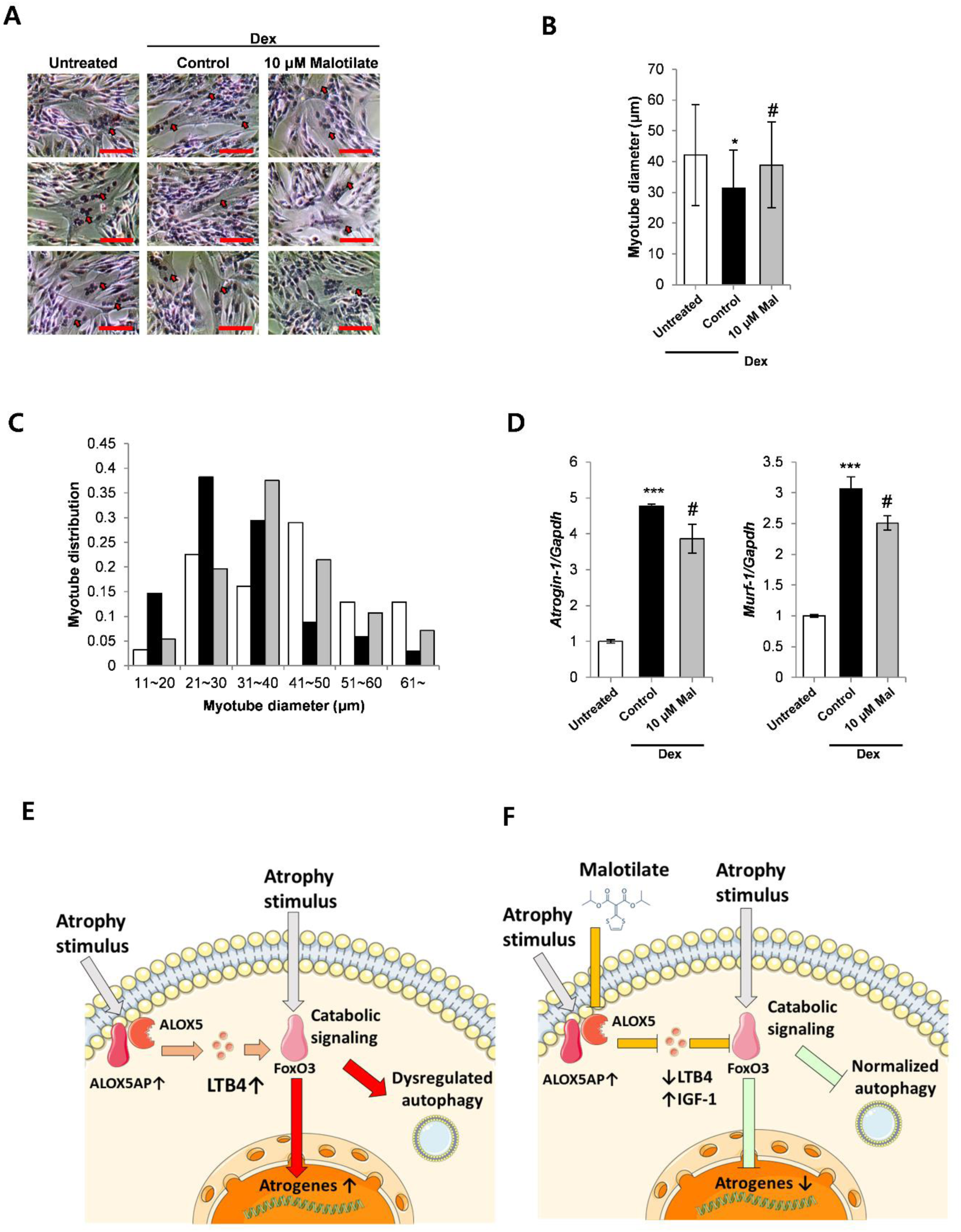
A) Representative H&E stained images of human donor myoblasts cultured as follows: (1) differentiation media (DM) for 72 h (untreated); (2) DM for 48 h and DM plus 10 μM Dex for 24 h; (3) DM for 48 h and DM plus 10 μM Dex and 10 μM malotilate (Mal) for 24 h (scale bar=100 μm). B) Mean myotube diameter. C) Myotube diameter distribution. D) qPCR analysis of atrogin-1 and MuRF-1 expression. E-F) Working model of the effect on malotilate on skeletal muscle atrophy. E) Under atrophy-inducing conditions, such as glucocorticoid treatment or aging, catabolic signaling pathways are activated in skeletal muscle fibers, producing atrophy effector mechanisms including upregulated atrogenes expression and dysregulated autophagy. Alox5 activating protein (Alox5AP) is also known to be upregulated under atrophy-inducing conditions, and the Alox5 product, LTB4, has been shown to increase the activity of catabolic regulators, such as FoxO signaling [29-31, 38-40], which would enhance muscle fiber atrophy. F) Malotilate treatment blocks the activity of Alox5, which would reduce cellular levels of LTB4 and upregulate the expression of anti-atrophy factors, such as IGF-1, inhibiting catabolic pathways. This could produce a beneficial effect against the effector pathways of skeletal muscle fiber atrophy.

## Discussion

Skeletal muscle atrophy can arise from a number of different conditions and it produces a major socioeconomic impact. For example, hospitalizations in individuals with aging-related muscle atrophy (termed sarcopenia) has been estimated to cost in excess of $19 billion in the United States [22]. Skeletal muscle atrophy arising from glucocorticoid use also is associated with a high personal and economic burden. Currently, there is no clinically approved drug for these forms of muscle atrophy [21]. In this study, we investigated the repurposing of malotilate, a clinically safe drug developed for liver regeneration and Alox5 inhibitor, as a muscle atrophy treatment. Our results indicate that malotilate is an attractive candidate for further development to treat muscle atrophy and Alox5 is a novel drug target for this disorder.

Malotilate treatment affected both the ubiquitin-proteasome proteolysis system (UPS) pathway and autophagy in myotubes undergoing atrophy. Malotilate inhibited the activation of Foxo3a, which is a major activator of the UPS. It has been reported that 75% of the protein degradation that occurs during skeletal muscle atrophy is contributed by the UPS [23]. Therefore, the protective effect of malotilate on muscle atrophy may be primary due to inhibition of Foxo3a. It has been reported that upon autophagy induction, for instance by nutrient starvation or glucocorticoids, LC3-I converts to LC3-II and induces a concurrent decrease in p62 [24]. Malotilate treatment normalized the ratio of LC3-I/LC3-II and increased expression of p62. It may be concluded that the anti-atrophy effects of malotilate treatment are mainly via UPS inhibition, along with a prevention of autophagy induction.

Malotilate did not significantly affect myoblast differentiation or the expression of key myogenic regulatory genes. Thus, the anti-atrophy effects of malotilate appear to be related to anti-atrophy mechanisms in differentiated muscle fibers, such as the inhibition of atrogene expression, rather than enhancing muscle stem cell differentiation. It should be noted that under conditions of muscle atrophy, such as aging, changes in the muscle stem cell niche are thought to be responsible for ineffective muscle regeneration, rather than abnormalities in stem cell proliferative responses.

Malotilate treatment increased skeletal muscle mass without affecting total body mass. Agents producing muscle atrophy, such as increased glucocorticoid signaling, are also known to increase adiposity [25]. The dual, opposing effects of malotilate on glucocorticoid signaling in muscle and adipose tissues may provide an explanation for the unchanged body mass observed in this study. The anti-muscle atrophy effect of malotilate was indicated to be due to the inhibition of Alox5, because gene knockdown of Alox5 also protected against glucocorticoid-induced atrophy. Although the inhibition of other members of the lipoxygenase family have been shown to influence skeletal muscle regeneration, the effect of malotilate is likely to be specific to Alox5, because malotilate has been shown to specifically target Alox5 compared to other members of the lipoxygenase family [18]. In the aging model, malotilate treatment significantly improved grip strength and also increased performance on the rotarod, although this was not statistically significant. Grip strength is used as the primary parameter of sarcopenia by the European Working Group on Sarcopenia in Older People (EWGSOP) [26]. These results suggest that malotilate is a candidate for preventing loss of muscle function and progression to sarcopenia, and should be further characterized in other sarcopenia models, such as the senescence-associated mouse or optic atrophy 1 knockout mouse [27].

The results in this study indicate that malotilate-mediated inhibition of Alox5 is a novel therapeutic approach for treating skeletal muscle atrophy. In addition, Alox5 gene knockdown prevented atrophy in myotubes and treatment with the Alox5 product, LTB4, induced atrophy. LTB4 levels were also found to be elevated in skeletal muscle undergoing atrophy. To our knowledge, there is no report showing that Alox5 is upregulated in skeletal muscle fibers undergoing atrophy. Alox5 activity is regulated by arachidonate 5-lipoxygenase-activating protein (Alox5AP, also known as 5-lipoxygenase activating protein, or FLAP), which anchors ALOX5 to the cell membrane and transfers the arachidonic acid substrate [28]. Interestingly, Alox5AP has been shown to be upregulated in aged human skeletal muscle and by glucocorticoid treatment [29]. The increased expression of Alox5AP under conditions of atrophy could explain the effects of malotilate observed in this study. In addition, the Alox5 product, LTB4, activates the BLT1/2 G-protein coupled signaling pathway and increases the activity of NF-κb signaling in atherosclerotic plaques and FoxO signaling in adipocytes [30, 31]. NF-κb and FoxO signaling are both master regulatory pathways of skeletal muscle atrophy [32]. Taken together, these previous findings and the results of this study support a mechanism of action wherein atrophy conditions upregulate Alox5AP expression, which enhances the activity of Alox5 and increases LTB4 levels. LTB4 stimulates the activity of atrophy signaling pathways, such as NF-κb and FoxO, to promote the effectors of myofiber catabolism, such as upregulated atrogenes expression and dysregulated autophagy. Treatment with malotilate reduces LTB4 levels and the activity of atrophy signaling pathways, producing an inhibition of muscle atrophy. A model of the effect of malotilate treatment and Alox5 inhibition on skeletal muscle atrophy is shown in Figure 8E-F.

Malotilate has been reported as a specific inhibitor of Alox5 [18]. The RNA seq data presented in this study indicates that Alox5 inhibition mediated by malotilate affects metabolic pathways (by KEGG analysis) and animal organ development (by GO analysis) in skeletal muscle cells. The primary metabolic changes associated with muscle atrophy caused by glucocorticoids, such as Dex, are insulin resistance, hyperglycemia, and hyperlipidemia [33]. Thus, targeting metabolic pathways using the Alox5 inhibitor malotilate has the potential to correct these changes in metabolism caused by Dex. The molecular mechanisms underlying organ development during embryogenesis have many similarities with the processes regulating tissue regeneration [34]. Therefore, the major effect on organ development indicated by GO analysis may explain the ability of this inhibitor to ameliorate the progression of skeletal muscle atrophy in the Dex treatment and aging animal models. Insulin-like growth factor-1 (IGF-1), a key modulator of muscle hypertrophy, was upregulated by malotilate treatment. IGF-1 levels are suppressed in many chronic diseases and produces muscle atrophy by the combined effects of suppressed protein synthesis, UPS activity, autophagy, and fiber regeneration [35]. Supplementation with IGF-1 alone has been shown to be effective for preventing muscle atrophy [36]. Interestingly, this upregulation of IGF-1 was specific for myotubes undergoing atrophy and not observed in normal myotubes treated with malotilate, indicating a disease-specific effect of drug treatment. Other differentially expressed genes that are known to influence muscle atrophy include upregulated superoxide dismutase, extracellular, which was previously shown to inhibit skeletal muscle wasting in heart failure, and downregulated signal transducer and activator of transcription 6 (STAT6), which is an inhibitory factor for adult myogenesis [37].

In summary, malotilate, a previously characterized liver regeneration drug studied in clinical trials, prevented skeletal muscle atrophy in multiple models and was effective using oral delivery. This drug also prevented atrophy in human skeletal myotubes. Malotilate treatment produced marked effects on muscle mass and performance in the Dex atrophy model, and the preservation of fast twitch fibers types in aged muscle, indicating that the Alox5 target may have a major effect on the progression of muscle atrophy. A role for Alox5 in muscle atrophy was also supported by studies of gene knockdown, supplementation with the LTB4 product, and assessment of LTB4 levels and Alox5 expression. In light of the clinical need to develop new drugs and targets for skeletal muscle atrophy, malotilate and Alox5 inhibitors can be attractive drug candidates for repurposing to treat this disorder.

## Acknowledgements

The authors of this manuscript certify that they comply with the ethical guidelines for authorship and publishing in the *Journal of Cachexia, Sarcopenia and Muscle* [38]. Funding: This work was supported by the National Research Foundation of Korea (NRF) funded by the Korean government (MSIT) (NRF-2020R1A2C2014194) and the Bio & Medical Technology Development Program of the National Research Foundation (NRF) funded by the Korean government (MSIT) (No. NRF-2020M3A9G3080282). This work was partly supported by the Institute for Information and Communications Technology Promotion (IITP) grant funded by the Korea government (MSIP; No. 2019-0-00567, Development of Intelligent SW systems for uncovering genetic variation and developing personalized medicine for cancer patients with unknown molecular genetic mechanisms), and a (GRI) ARI” grant fund in 2022. The funders had no role in study design, data collection and analysis, decision to publish, or preparation of the manuscript. Figure 7D was produced under a Creative Commons Attribution 3.0 Unported License using https://smart.servier.com/. Author contributions: H.K. carried out experiments, analyzed the data and wrote the manuscript. S-H. L. and S-W. K. carried out experiments. D-W. J. and D.W. acquired funding, supervised the research, and wrote the manuscript. Conflict of interest: H.K., D.-W.J. and D.R.W. are named co-inventors of a pending provisional patent application based in part on the research reported in this paper.

---

**Supplementary Figure 1:**
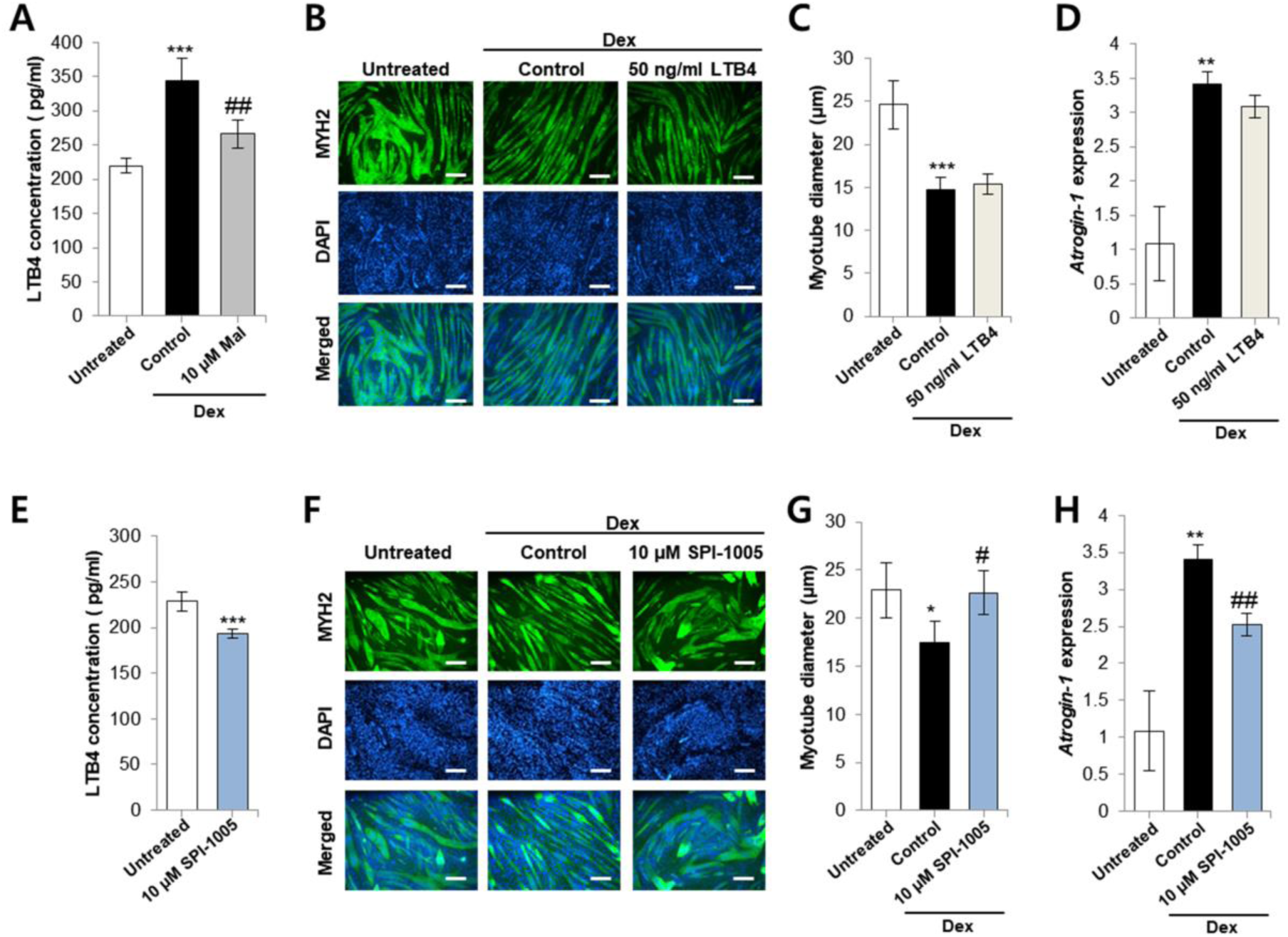
A) ELISA analysis LTB4 levels in C2C12 myoblasts cultured as follows: 1) 120 h incubation with DM; (2) 96 h incubation with DM, and 24 hour treatment with 10 μM Dex; (3) 96 h incubation with DM, and 24 hour treatment with 10 μM Dex and 10 μM malotilate (Mal). B) MYH2 immunocytochemistry of C2C12 myoblasts cultured as follows: (1) 120 h incubation with DM plus control; (2) 96 h incubation with DM, and 24 h treatment with 10 μM Dex; (3) 96 h incubation with DM, and 24 h treatment with 10 μM Dex plus 50 ng/mL LTB4 (scale bar=100 μm). C) Myotube diameter. D) qPCR analysis of atrogin-1 expression in C2C12 myoblasts cultured as described in part (B). E) ELISA analysis of LTB4 levels in C2C12 myoblasts cultured as follows: 1) 120 h incubation with DM; (2) 96 h incubation with DM, and 24 h treatment with 10 μM SPI-1005; F) MYH2 immunocytochemistry of C2C12 myoblasts cultured as follows: (1) 120 h incubation with DM; (2) 96 h incubation with DM, and 24 h treatment with 10 μM Dex; (3) 96 h incubation with DM, and 24 h treatment with 10 μM Dex plus 10 μM SPI-1005 (scale bar=100 μm). G) Myotube diameter. H) qPCR analysis of atrogin-1 expression in C2C12 myoblasts cultured as described in part (F). *=*p*<0.05, **=*p*<0.01, ***=*p*<0.001 compared to untreated. ^#^=*p*<0.05, ^##^=*p*<0.01 compared to Dex treatment alone.

**Supplementary Figure 2:**
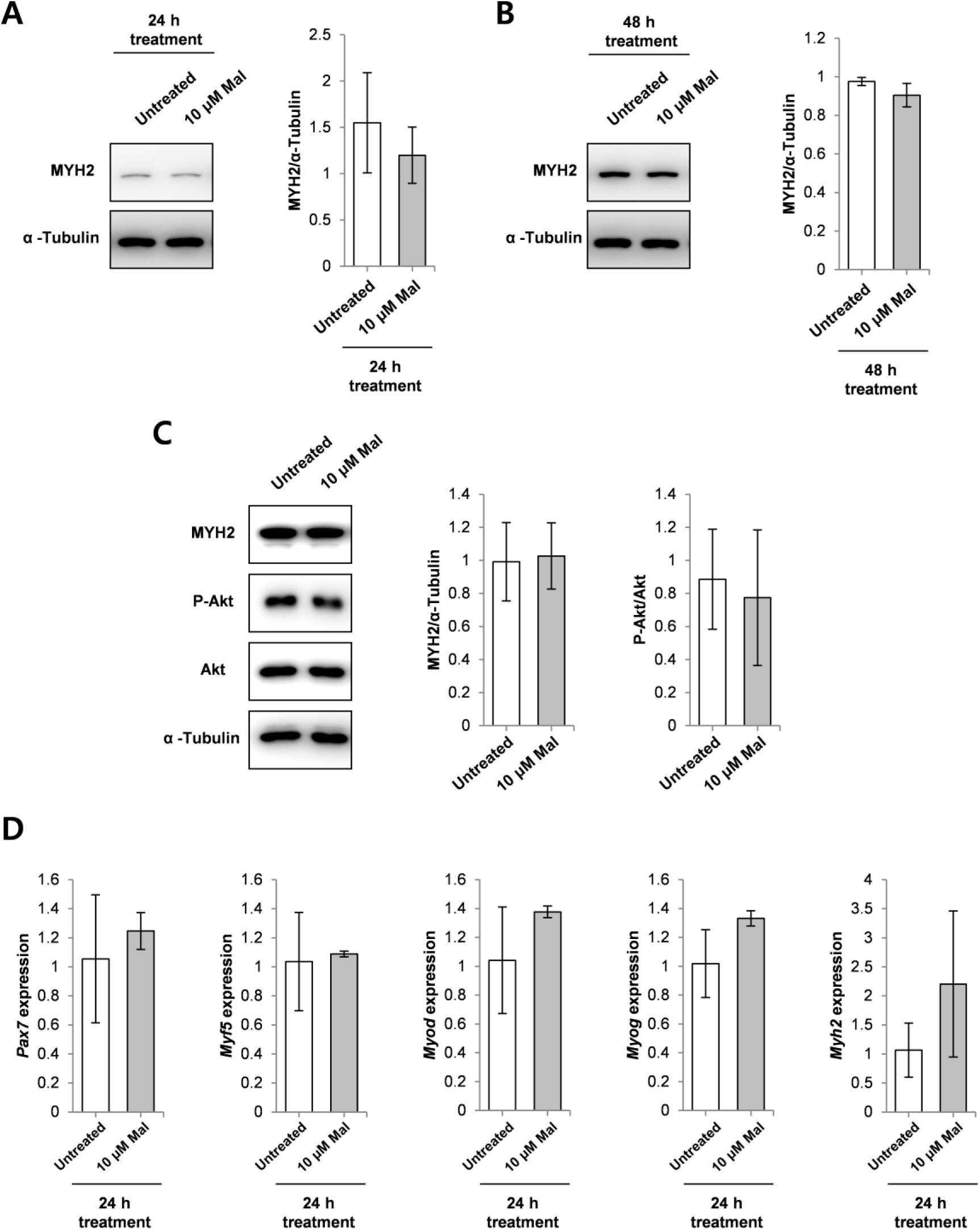
A-B) Western blot analysis of Myh2 expression in C2C12 myoblasts cultured with DM or DM plus 10 μM malotilate (Mal) for 24-48 h. α-Tubulin was used as a loading control. C) Western blot analysis of Akt phosphorylation, Akt expression, and Myh2 expression in in C2C12 myoblasts cultured as follows (1) 120 h incubation with DM; (2) 96 h incubation with DM, and 24 h treatment of 10 μM Malotilate; α-Tubulin was used as a loading control. Quantification of Myh2 expression and Akt phosphorylation are shown. D) qPCR analysis of Pax7, Myf5, MyoD, MyoG, and Myh2 in C2C12 myoblasts cultured with DM or DM plus 10 μM malotilate for 24 h.

**Supplementary Figure 3:**
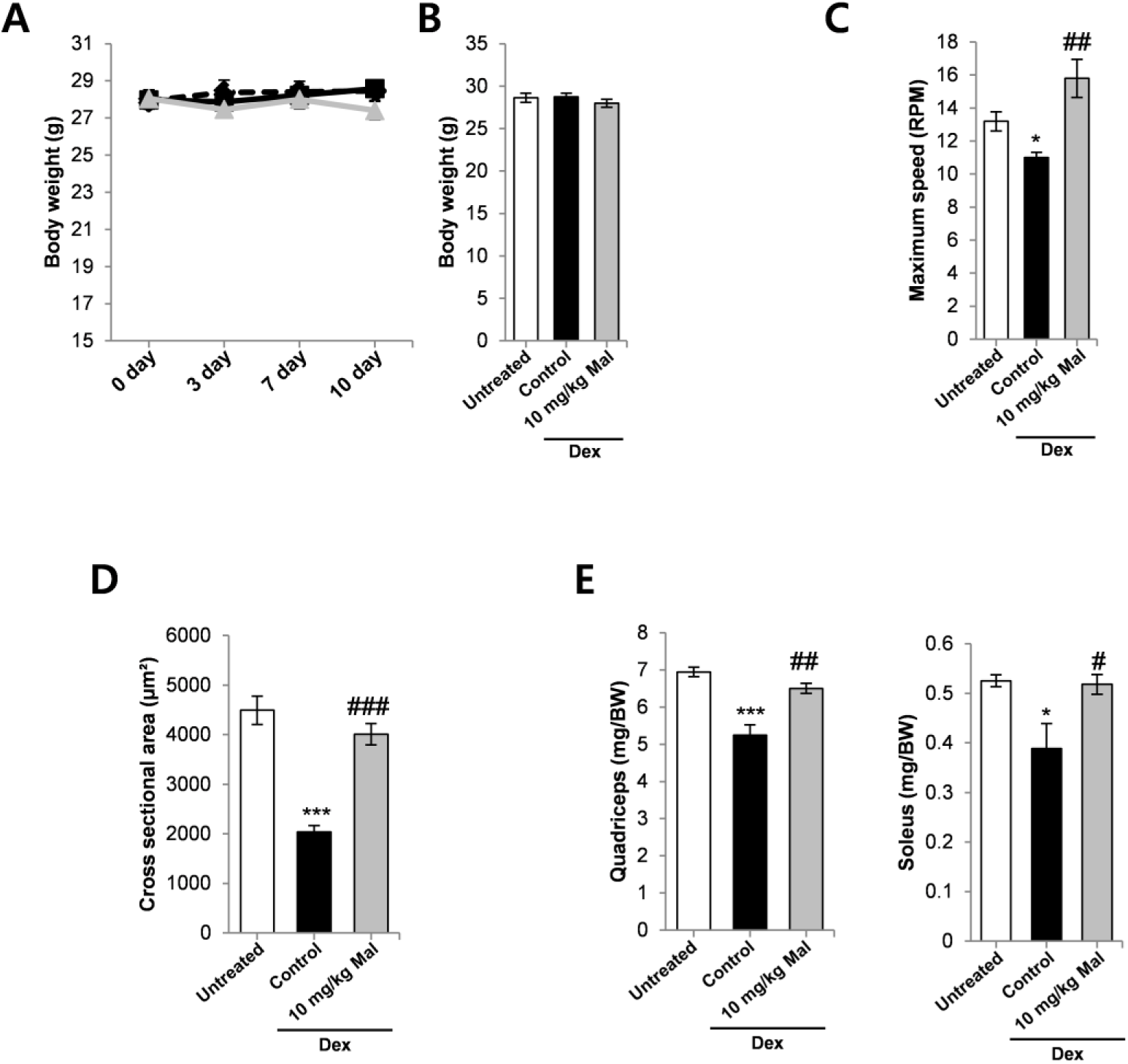
A) Body weight in mice treated with vehicle alone, Dex, or Dex plus malotilate (Mal) during 14 d treatment. B) Final body weight. C) Maximum speed achieved on the rotarod. D) Fiber cross sectional area and distribution of the quadriceps. E-F) Quadriceps and soleus mass. *=*p*<0.05, ***=*p*<0.001 compared to vehicle treated mice. ^#^=*p*<0.05, ^##^=*p*<0.01 compared to Dex treated mice (n=4-5 per experimental group).

**Supplementary Figure 4:**
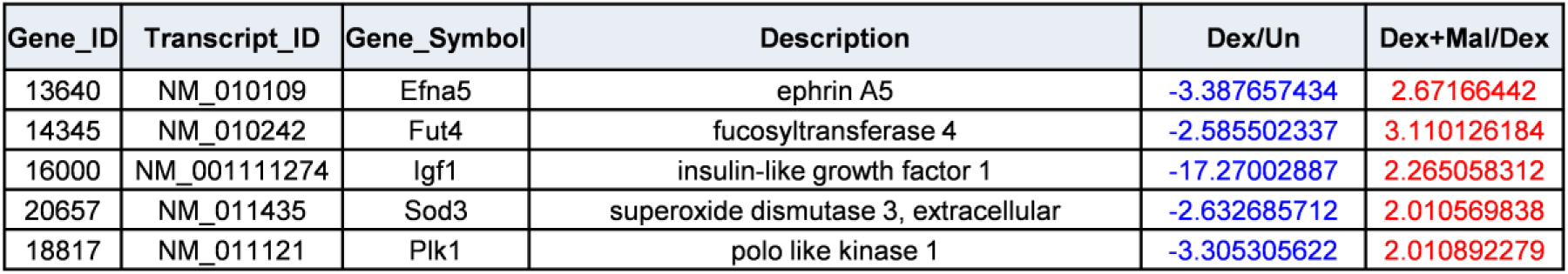
Related to Figure 7F: Selected genes known to be implicated in skeletal muscle atrophy showing differential expression in Dex and Dex_Mal, compared to Dex and Un.

**Supplementary Table 1.** Primaiy antibodies used in this study

| **Primary antibody** | **Clone** | **Company** | **Catalog No.** | **Dilution** |
| --- | --- | --- | --- | --- |
| α-Tubulin | Polyclonal | Invitrogen | PA5-29444 | 1:5000 |
| Puromycin (12D10) | Monoclonal | Millipore | MABE-343 | 1:25000 |
| Atrogin-1/MAFbx (F-9) | Monoclonal | SANTA CRUZ | Sc-166806 | 1:100 |
| FoxO3a (D19A7) | Monoclonal | CST | #12829 | 1:1000 |
| Phospho-FoxO3a (Ser253) | Polyclonal | CST | #9466 | 1:1000 |
| Myosin heavy chain 2 (A4.7) | Monoclonal | SANTA CRUZ | Sc-53095 | 1:1000 |
| P62/SQSTM1 | Polyclonal | Merck | P0067 | 1:1000 |
| LC3B | Polyclonal | Merck | L7543 | 1:1000 |
| 5-Lipoxygenase | Monoclonal | Abcam | AB169755 | 1:1000 |
| Phospho-Akt (Ser473) | Polyclonal | CST | #9271 | 1:1000 |
| Akt | Polyclonal | CST | #9272 | 1:1000 |
| Laminin | Polyclonal | Abcam | AB11575 | 1:30 |
| Myosin heavy chain Type IIA | Monoclonal | DSHB | SC-71 | 2-5 ug/ml |
| Myosin heavy chain Type IIB | Monoclonal | DSHB | BF-F3 | 2-5 ug/ml |

**Supplementary Table 2.** Secondaiy antibodies used in this study

| **Secondary antibody** | **Conjugated use** | **Company** | **Catalog No.** | **Dilution** |
| --- | --- | --- | --- | --- |
| Horse anti-Mouse IgG<br>HRP | HRP | CST | #7074 | 1:2000 |
| Goat anti-Rabbit IgG<br>HRP | HRP | CST | #7074 | 1:2000 |
| Alexa Fluor™ 488<br>Goat anti-mouse<br>IgG(H+L) | Alexa Fluor<br>488 | INVITROGEN | A11001 | 1:1000 |
| Goat anti-Rabbit IgG<br>h+l Dylight® 594<br>Conjugated | Dylight®<br>594 | Bethyl | A90-116D4 | 1:1000 |

**Supplementary Table 3.**
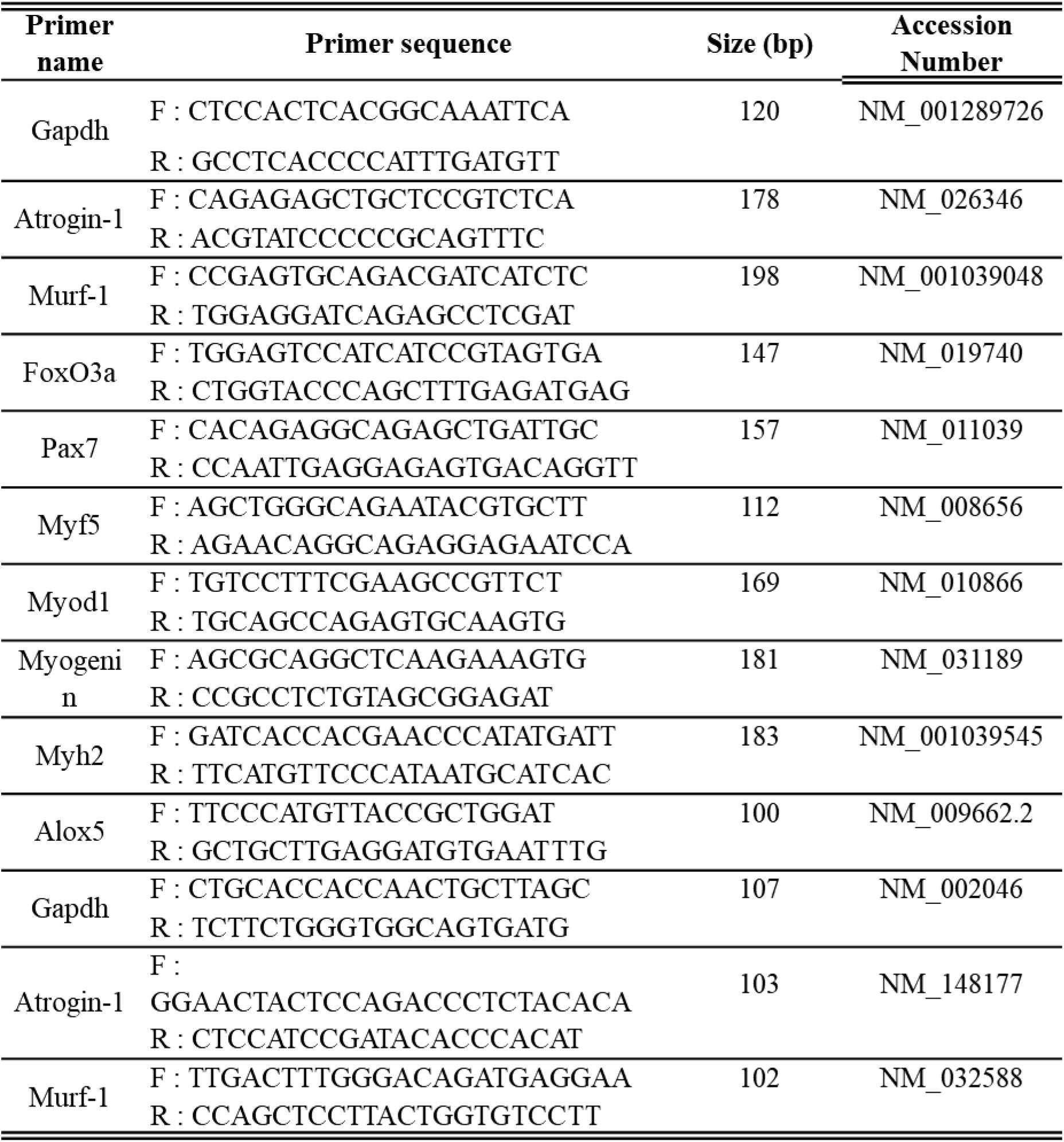
Primers used for qPCR

